# NCOA4 initiates ferritinophagy by binding GATE16 using two highly avid short linear interaction motifs

**DOI:** 10.1101/2024.06.09.597909

**Authors:** April Lee, Joseph H. Davis

## Abstract

Cells carefully regulate cytosolic iron, which is a vital enzymatic cofactor, yet is toxic in excess. In mammalian cells, surplus iron is sequestered in ferritin cages that, in iron limiting conditions, are degraded through the selective autophagy pathway ferritinophagy to liberate free iron. Prior work identified the ferritinophagy receptor protein NCOA4, which links ferritin and LC3/GABARAP-family member GATE16, effectively tethering ferritin to the autophagic machinery. Here, we elucidate the molecular mechanism underlying this interaction, discovering two short linear motifs in NCOA4 that each bind GATE16 with weak affinity. These binding motifs are highly avid and, in concert, support high-affinity NCOA4•GATE16 complex formation. We further find the minimal NCOA4^383-522^ fragment bearing these motifs is sufficient for ferritinophagy and that both motifs are necessary for this activity. This work suggests a general mechanism wherein selective autophagy receptors can distinguish between the inactive soluble pools of LC3/GABARAPs and the active membrane-conjugated forms that drive autophagy. Finally, we find that iron decreases NCOA4^383-522^’s affinity for GATE16, providing a plausible mechanism for iron-dependent regulation of ferritinophagy.

## INTRODUCTION

All cells must tightly regulate nutrient levels, including metals such as iron that act as crucial cofactors for essential enzymes. Whereas ample labile iron is vital for cellular function, excess iron can be toxic due, in part, to its ability to form damaging reactive oxygen species (Wang & Pantopoulos, 2011). Indeed, iron dysregulation has been connected to a variety of neurological diseases including hereditary ferritinopathy, Alzheimer’s (Ward et al., 2014) and Parkinson’s (Sofic et al., 1988) diseases. Mammalian cells are thought to maintain an appropriate pool of cytosolic iron by balancing iron uptake and storage in ferritin cages with the release of ferritin-sequestered iron through degradation of these proteinaceous cages (Arosio et al., 2009).

One mechanism by which ferritin is degraded upon depletion of free iron pools is ferritinophagy, a type of selective autophagy (Mancias et al., 2015) wherein select cellular components are recognized by an autophagy receptor protein and encapsulated in a double-membraned vesicle that is targeted to the lysosome for proteolysis (Zaffagnini & Martens, 2016). Ferritinophagy is thought to rely on the protein Nuclear Receptor Coactivator 4 (NCOA4), which was first identified as a selective autophagy receptor in an LC-MS/MS based proteomics approach targeting proteins that copurified with autophagosomes (Mancias et al., 2014). The activity of NCOA4 is regulated via iron-dependent (Zhao et al., 2024) changes in affinity for ferritin as well as an E3 ligase HERC2, which can promote NCOA4 degradation via the proteasome (Mancias et al., 2015). An NCOA4 fragment composed of residues 383-522 (NCOA4^383-522^, hereafter) was subsequently found to be sufficient to bind directly to ferritin and to interact with proteins in the autophagic machinery (Hoelzgen et al., 2024; Mancias et al., 2015).

To link the intended substrate and the autophagic machinery, selective cargo receptors typically bind to one or more of the six members of the Microtubule-associated proteins 1A/1B light chain 3B (LC3/GABARAP) family of integral autophagosomal proteins (Lee & Lee, 2016). These homologs, which include LC3A, LC3B, LC3C, GABARAP, GEC1, and GATE16, exchange between a monomeric soluble form, and a lipid-conjugated form wherein they are covalently linked to phosphatidyl ethanolamine headgroups of phospholipids in the autophagosomal membrane (Kabeya et al., 2004; Kraft et al., 2014). Selective autophagy receptors are thought to tether bound substrates to the autophagosomal membrane by simultaneously binding to LC3/GABARAPs via a short linear motif known as an LC3 interacting region (LIR). These motifs are hydrophobic in nature and have a consensus sequence of [W/F/Y]-X-X-[I/L/V], often accompanied by an N-terminal patch of acidic residues (Johansen & Lamark, 2020). Structural studies have further shown that the LC3/GABARAPs contain a LIR docking site (LDS) consisting of two hydrophobic pockets that support docking of the aromatic and aliphatic residues of the LIR motif (Johansen & Lamark, 2020).

Autophagy receptors are typically thought to associate with LC3/GAPARAPs using a single LIR, with affinity often in the low micromolar range (Johansen & Lamark, 2020). However, the LIR motif is common in the proteome and multiple instances can be found in most autophagy receptors, raising the possibility of highly avid, multivalent interactions to arrays of membrane tethered LC3/GABARAP-family proteins. In such a model, multiple binding motifs on the same receptor, each with weak intrinsic affinity, would act in concert to increase the effector receptor protein concentration and thus support high affinity binding to multimerized LC3/GABARAPs (Errington et al., 2019; Lluís Garcés et al., 2009) such as those found on an autophagosomal membrane densely decorated by LC3/GABARAPs.

Here, to biochemically define the interactions supporting ferritinophagy, we extend that work of Mancias et al., who previously identified the LC3/GABARAP family protein GATE16 as a strong NCOA4 interactor (Mancias et al., 2014). Specifically, using a purified NCOA4^383-522^ fragment, we identify and characterize two LIR-like motifs in NCOA4^383-522^ that each weakly bind to GATE16. We further show that these LIR-like motifs are highly avid and that robust GATE16•NCOA4 complex formation requires oligomerized GATE16. Furthermore, we show that this minimal NCOA4^383-522^ fragment is sufficient for lysosomal degradation of ferritin in cells and that such degradation requires the two identified motifs. Finally, we demonstrate that binding of iron to NCOA4^383-522^ decreases its affinity for GATE16.

## RESULTS

### NCOA4 binds directly to GATE16

Though it has been previously reported that NCOA4 interacts with GATE16, this interaction was exclusively demonstrated through co-immunoprecipitations from cell lysates and cellular co-localization studies (Dowdle et al., 2014; Mancias et al., 2014). Such evidence does not preclude indirect GATE16•NCOA4 binding through a protein intermediate. Thus, we sought to determine whether NCOA4 and GATE16 can, in fact, bind directly. For these experiments we used the biochemically amenable fragment of NCOA4^383-522^ (**Figure 1a-b**) that is sufficient to bind ferritin (Hoelzgen et al., 2024), but lacks both the putative coiled-coil domain and domains that overlap with the β-isoform, which does not bind ferritin or support ferritinophagy (Mancias et al., 2015). After purifying NCOA4^383-522^ and GST-GATE16 which, in our assay, were monomeric and dimeric, respectively (**Supplementary Figure 1**), we assessed binding via far-western blots and biolayer interferometry (BLI). We observed concentration-dependent association in both assays (**Figure 1c-d**) consistent with NCOA4^383-522^ having directly bound to this dimeric (GST-GATE16)_2_. Notably, careful inspection of the BLI traces revealed biphasic binding kinetics in the association and dissociation phases, consistent with the presence of at least two binding events (**Supplementary Figure 2**).

**Figure 1.**
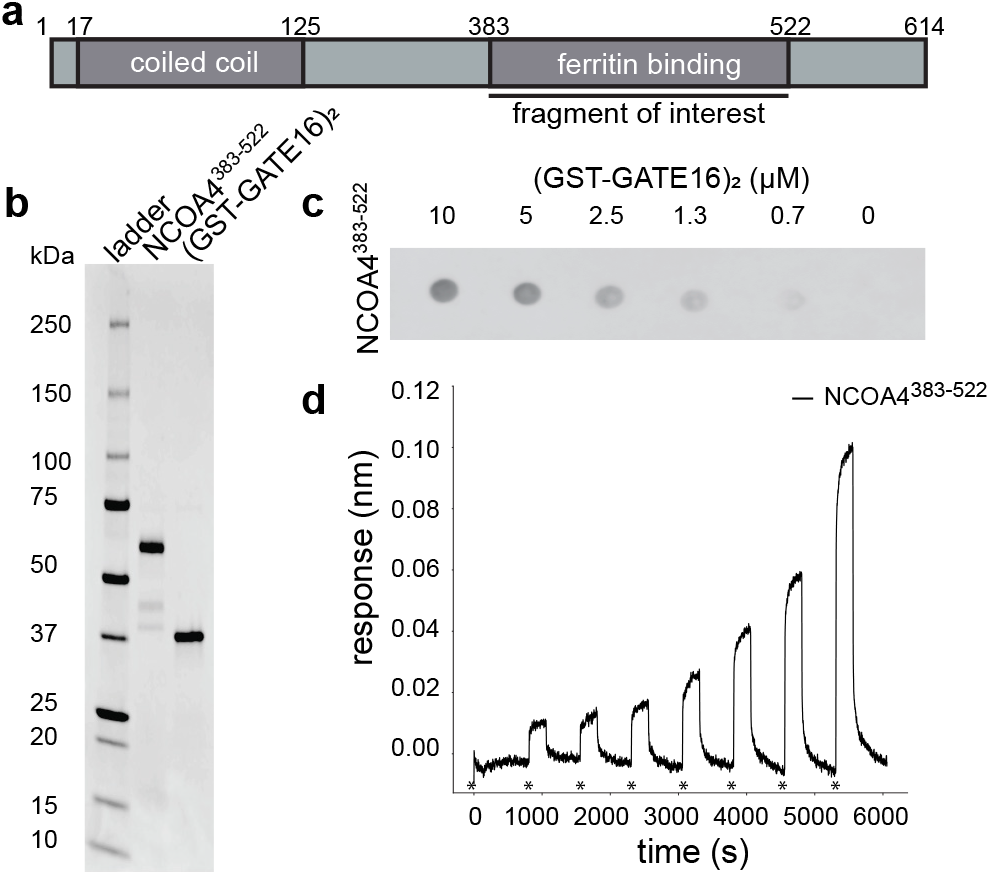
NCOA4^383-522^ binds directly to (GST-GATE16)_2_. **(a)**Schematic of NCOA4 domain architecture. The fragment of interest (NCOA4^383-522^) used here is noted. **(b)** SDS-PAGE gel of purified biotinylated AviTag-His_6_-NCOA4^383-522^-MBP (NCOA4^383-522^) and (GST-GATE16)_2_. **(C)**Far-western of NCOA4^383-522^ binding to decreasing concentrations of (GST-GATE16)_2_ which was spotted on the membrane at indicated concentration. Signal was detected by HRP-conjugated streptavidin directed to soluble, Avi-tagged NCOA4^383-522^, which was incubated on membrane at a concentration of 0.1μM. **(d)** Biolayer interferometry (BLI) trace of NCOA4^383-522^ binding to (GST-GATE16)_2_ where NCOA4^383-522^ was immobilized to the probe and was incubated with increasing concentrations (0nM, 78nM, 156nM, 313nM, 625nM, 1.25μM, 2.5μM, 5μM) of (GST-GATE16)_2_. Initialization of each incubation noted by asterisks. Trace depicted is background corrected for binding of (GST-GATE16)_2_ alone to the probe at each concentration.

### NCOA4^383-522^ binds to GATE16 via two LIR-like motifs

NCOA4^383-522^ contains several putative LIR-like sequence motifs. To determine if such motifs supported GATE16•NCOA4 binding, we constructed a tiled array of peptides, each 20 amino acids long, that spanned the NCOA4^383-522^ sequence, and we probed this array for binding with (GST-GATE16)_2_. The peptide array revealed two candidate linear interacting regions within the NCOA4 fragment – specifically ^413^FAECV^417^ and ^485^SFQVI^489^ (**Supplementary Figure 4**). Of note, whereas the ^485^S<u>F</u>QVI^489^ motif contained a canonical LIR sequence, the ^413^<u>F</u>AE<u>C</u>V^417^ motif bore cysteine substituted for the L/I/V residue specified by the canonical four residue LIR motif. We found that these identified motifs were highly conserved amongst NCOA4 orthologs, with the limited observed sequence variation preserving the hydrophobic nature of the key residues, including this noted cysteine (**Supplementary Figure 5**). To directly assess the impact of these motifs on NCOA4•GATE16 binding, we individually substituted alanine for the key aromatic residues (^413^F:A; ^486^F:A) in the identified LIR motifs in NCOA4^383-522^ and tested the impact of these mutations on (GST-GATE16)_2_ binding using both far-western blots (**Figure 2a**) and BLI (**Figure 2b**). In each assay, mutations to these putative binding motifs decreased (GST-GATE16)_2_ binding, consistent with the necessity of each motif. Notably, the effect was not due to simply unfolding NCOA4^383-522^, as the mutations did not impact global protein stability as assayed by a SYPRO Orange protein thermal stability shift assay (**Supplementary Figure 6**).

**Figure 2.**
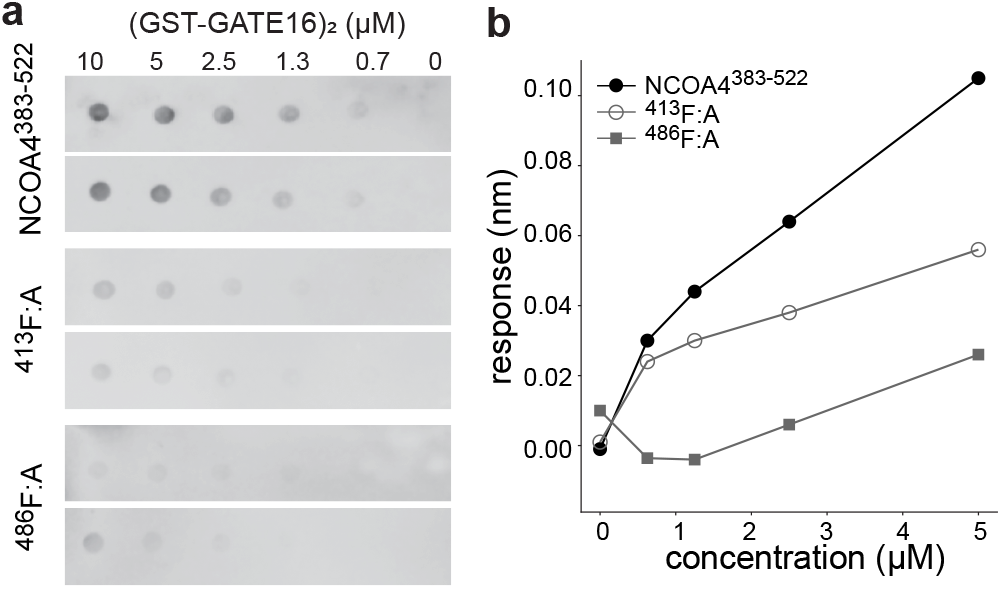
NCOA4^383-522^ binds to (GST-GATE16)_2_ through two LIR-like motifs. **(a)**Far-western of NCOA4^383-522^ and NCOA4^383-522^ motif mutants binding to (GST-GATE16)_2_, which was spotted on membrane at indicated concentrations. Binding was detected using HRP-conjugated streptavidin directed to soluble, Avi-tagged NCOA4^383-522^, which was incubated on membrane at a concentration of 0.1μM. NCOA4^383-522^ and motif mutants assayed in duplicate. **(b)**BLI response curve of NCOA4^383-522^ and motif mutants binding to (GST-GATE16)_2_, where the NCOA4^383-522^ construct was immobilized to the probe and tested against increasing concentrations (0nM, 625nM, 1.25μM, 2.5μM, 5μM) of (GST-GATE16)_2_. FAECV motif mutant noted as ^413^F:A and S<u>F</u>QVI mutant labeled as ^486^F:A.

### NCOA4•GATE16 binding relies on avidity *in vitro*

Having identified two motifs in NCOA4^383-522^ that were each necessary for GATE16 association, we next sought to determine where on GATE16 those peptides bound. Since LIR motifs typically bind to LC3/GABARAP family proteins in regions known as LIR-docking sites (LDS) (Johansen & Lamark, 2020), we hypothesized the LDS was a likely binding location. To assess this, we performed competitive fluorescence anisotropy assays in which an unlabeled peptide containing either of the identified NCOA4^383-522^ motifs was added to a solution bearing monomeric GATE16 and a fluorescently labeled peptide derived from ATG4B that is known to bind to the LDS (Skytte Rasmussen et al., 2017). Though the NCOA4^383-522^-derived peptides displaced the ATG4B peptide in a sequence-dependent manner, consistent with competitive binding to the LDS, they did so with weak apparent affinity (**Figure 3a-b**).

**Figure 3.**
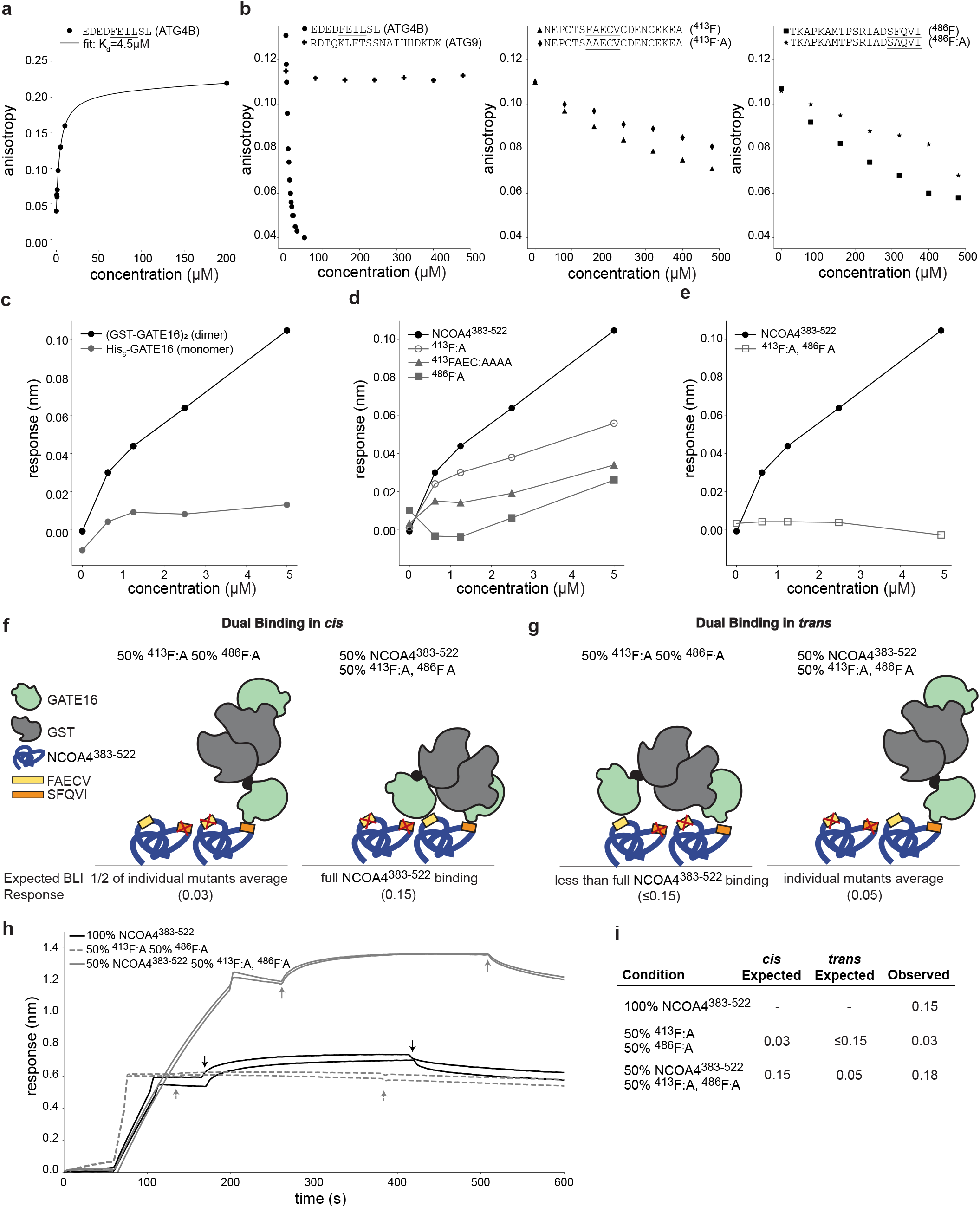
Binding of NCOA4^383-522^ to (GST-GATE16)_2_ requires avidity. **(a)**Fluorescence anisotropy of dye-labeled peptide containing the ATG4B LIR binding to His_6_-GATE16. Data fit to standard binding isotherm with apparent K_D_ noted. **(b)**Competition fluorescence anisotropy assay of dye-labeled ATG4B LIR peptide competing for binding to monomeric GATE16 against increasing concentrations of unlabeled peptides derived from ATG4B, ATG9, NCOA4^383-522^ or NCOA4^383-522^ motif mutants, as noted. **(c)**BLI response curve of NCOA4^383-522^ binding to His_6_-GATE16 or (GST-GATE16)_2_ where NCOA4^383-522^ construct is immobilized to the probe and tested against increasing concentrations (0nM, 625nM, 1.25μM, 2.5μM, 5μM) of either GATE16 construct. **(d,e)** BLI assay of (GST-GATE16)_2_ binding to noted NCOA4^383-522^ constructs, probed at (GST-GATE16)_2_ concentrations as described in (c). **(f,g)** Schematic of hypothesized binding modes for (GST-GATE16)_2_ binding in “*cis*” (f) or in “*trans*” (g) to NCOA4^383-522^. BLI response expected for each model at the noted ratios of each protein construct detailed below schematic. **(h)** BLI traces of assays performed as depicted in (f,g). Two replicates of each assay shown. Each assay used 5μM (GST-GATE16)_2_. **(i)** Expected responses for *cis* and *trans* models and quantification of response for BLI traces in (g).

The observed discrepancy in NCOA4 affinity for monomeric GATE16 and dimeric (GST-GATE16)_2_ led us to hypothesize that the two LIR motifs synergistically facilitated NCOA4•GATE16 binding. In our model, we reasoned that binding of one LIR-like motif to a dimerized copy of GATE16 could increase the effective concentration of the second LIR-like motif, thereby facilitating its binding to the GST-dimerized GATE16 copy. To more directly test this model, we assayed the binding of NCOA4^383-522^ to GATE16 as a function of GATE16’s oligomeric state using either monomeric His_6_-GATE16 or dimeric (GST-GATE16)_2_ (**Supplementary Figure 1**). Our BLI-based binding assay showed improved binding with (GST-GATE16)_2_ relative to monomeric His_6_-GATE16 (**Figure 3c**). Of note, GST alone was unable to bind NCOA4^383-522^ in any of our BLI, far-western, or peptide array assays (**Supplementary Figures 3-4**,**7**).

To further assess the impact of multiple NCOA4^383-522^ motifs on GATE16 binding, we generated single and double mutants of the identified motifs. Mutation of either motif alone significantly decreased binding to (GST-GATE16)_2_ as assayed by BLI, though neither was sufficient to completely abrogate binding (**Figure 3d**). Notably, replacing the FAECV motif with a tetra-alanine linker further abrogated binding relative to the ^413^F:A mutant, likely due to contribution of the cysteine (**Figure 3d**). In the double mutant (^413^F:A, ^486^F:A), we observed even weaker binding, consistent with each motif contributing to the overall NCOA4^383-522^•GATE16 interaction (**Figure 3e**).

We next assessed the oligomeric state of NCOA4^383-522^ in the context of our observed NCOA4^383-522^•GATE16 binding to obtain a more complete binding model. In contrast to previous studies reporting NCOA4^383-522^ can dimerize despite lacking the N-terminal coiled-coil oligomerization domain contained in the full-length protein (Gryzik et al., 2017), we found that NCOA4^383-522^ was monomeric at our working concentrations, as assessed by SEC-MALS (**Supplementary Figure 1**). However, since the BLI binding assays require that NCOA4^383-522^ be immobilized on an assay probe tip, it was still formally possible that NCOA4^383-522^ was positioned in such a way that it was acting as an artificial dimer. To assess whether NCOA4^383-522^ was acting as a monomer or a dimer in this assay, we performed two complementary tests via BLI, probing for NCOA4^383-522^ binding to (GST-GATE16)_2_ either in “*cis*” to monomeric NCOA4^383-522^, or in “*trans*” to an apparently dimeric form (**Figure 3f-g**). In the first assay, the two individual motif mutants (^413^F:A and ^486^F:A) were mixed in a 1:1 ratio, loaded on the probe and their binding to (GST-GATE16)_2_ was measured. If NCOA4^383-522^ were acting as a dimer in this assay, we expected binding similar to NCOA4^383-522^ lacking motif mutants, as each singly mutated NCOA4^383-522^ would still have one motif available to bind to one subunit of the GST-GATE16 dimer. Instead, we observed strongly reduced binding (**Figure 3h-i**). In a complementary assay, we mixed NCOA4^383-522^ and the double mutant (^413^F:A, ^486^F:A) in a 1:1 ratio, loaded the probe at 2-fold higher density and measured binding to (GST-GATE16)_2_. Of note, loading the probe with twice the amount of NCOA4^383-522^ produced a proportional increase in observed binding response (**Supplementary Figure 8**). As such, if NCOA4^383-522^ were acting as a monomer in this assay, then we would expect this protein mixture loaded to at the 2-fold higher density to produce similar binding to that observed with mutation-free protein loaded at the standard density. We observe binding at a level predicted by the monomeric model (**Figure 3f**) and thus conclude that a monomeric NCOA4^383-522^ uses two binding motifs to associate with dimeric GST-GATE16 (**Figure 3h-i**).

### NCOA4^383-522^ requires two LIR-like motifs to direct lysosomal degradation of ferritin *in vivo*

Having established the importance of these two LIR-like motifs *in vitro*, we next assessed the role of these motifs *in vivo*. To do so, we stably integrated NCOA4^383-522^ or variants mutated in the LIR-like motifs into NCOA4 knockout HeLa cells (NCOA4Δ, hereafter) (Gryzik et al., 2021). Of note, we measured similar levels of NCOA4 in each transfected cell line (**Supplementary Figure 9**). With these cell lines, we first determined whether the NCOA4^383-522^ fragment could facilitate the degradation of ferritin by comparing the extent of ferritin degradation in WT, NCOA4Δ and NCOA4^383-522^ cells upon iron chelation with deferoxamine (DFO). Using quantitative western blots (see Methods), we observed that ferritin was not degraded in NCOA4Δ cells, whereas ferritin degradation was robust in wild-type cells and those bearing the NCOA4^383-522^ (**Figure 4a,c**). Notably, the WT cells exhibited greater chelation-induced degradation than those expressing the NCOA4^383-522^ fragment, suggesting additional contributions to ferritinophagy outside of this biochemically amenable NCOA4 fragment.

**Figure 4.**
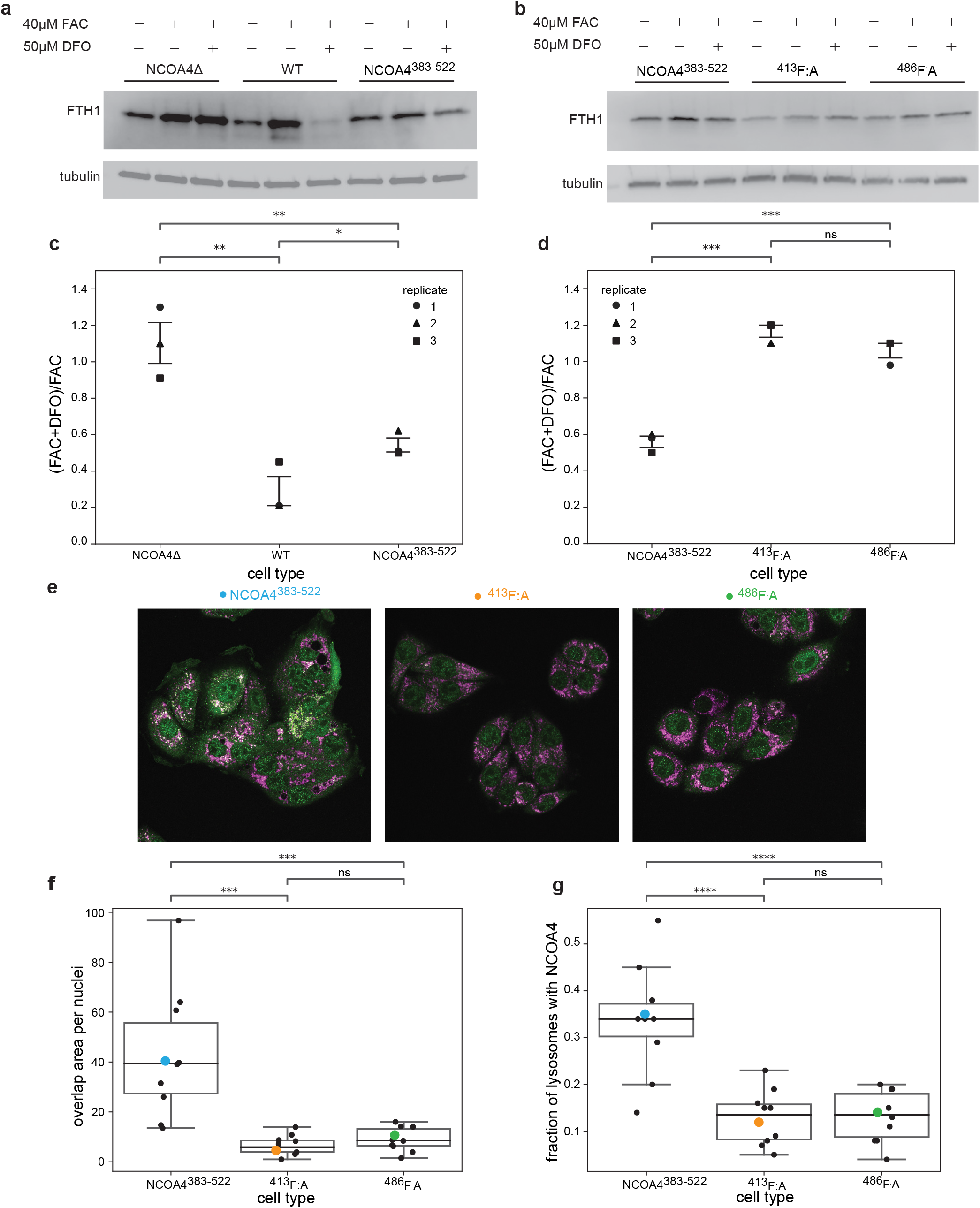
NCOA4^383-522^ supports ferritin degradation *in vivo* and requires both LIR-like motifs. **(a)**Western blots against ferritin heavy chain (FTH1) and tubulin in NCOA4Δ, WT and NCOA4^383-522^ cell lines. Cells either untreated, treated for 24 hours with ferric ammonium citrate (FAC), or treated with FAC before undergoing a 24-hour treatment with deferoxamine (DFO). **(b)**Western blots as described in (a) in either NCOA4^383-522, 413^F:A, or ^486^F:A cell lines.

Given that NCOA4383-522 facilitated ferritin degradation, we next assessed the impact of mutations to the two LIR-like motifs. Consistent with our *in vitro* binding data, mutation of either motif was sufficient to prevent NCOA4^383-522^-dependent ferritin degradation (**Figure 4b,d**), supporting the hypothesis that both motifs are required for stable NCOA4^383-522^ binding to LC3/GABARAP-family proteins and by extension for ferritin degradation in cells.

Finally, to assess whether the observed ferritin degradation occurred in the lysosome, we performed colocalization studies via immunofluorescence to measure the impact of these identified LIR motifs on NCOA4:lysosome colocalization. Specifically, we stained for GFP-NCOA4^383-522^ and lysosomes (LAMP1) in our NCOA4^383-522^ and NCOA4^383-522^ mutant cell lines in the presence of DFO. Strikingly, we saw significant colocalization of NCOA4^383-522^ with lysosomes, consistent with NCOA4-mediated ferritin degradation through the autophagy-lysosomal pathway (**Figure 4e**). Furthermore, when comparing the degree of colocalization in NCOA4^383-522^ versus the NCOA4^383-522^ motif mutant cell lines, we found that mutation of either NCOA4^383-522^ LIR-like motif resulted in significantly decreased NCOA4:lysosome colocalization (**Figure 4f,g**).

### Iron regulates the binding of NCOA4^383-522^ to GATE16

Given that the NCOA4•GATE16 interaction is crucial in the progression of ferritinophagy, an iron sensing pathway, we next asked whether iron could regulate this interaction. It was recently reported that NCOA4^383-522^ can bind an iron-sulfur cluster coordinated by four cysteines (Zhao et al., 2024). Importantly, one of these four cysteines, residue 416, is present in the ^413^FAE<u>C</u>V^417^ LIR-like motif we have described. Thus, we hypothesized that were NCOA4^383-522^ iron-loaded, the FAECV motif would be occluded, which would prevent robust GATE16 binding. To test this, we chelated iron from purified NCOA4^383-522^, or reconstituted the protein with iron anaerobically (see Methods). We used a ferene assay (Fish, 1988; Levitz et al., 2022; McCarthy & Booker, 2018) to measure the equivalents of iron bound in each sample (**Figure 5a**). For each of these samples we then measured binding to (GST-GATE16)_2_ and observed a marked decrease in (GST-GATE16)_2_ binding in the iron-bound NCOA4^383-522^ sample (**Figure 5b**), consistent with a role for iron in negatively regulating NCOA4•GATE16 complex formation.

**Figure 5.**
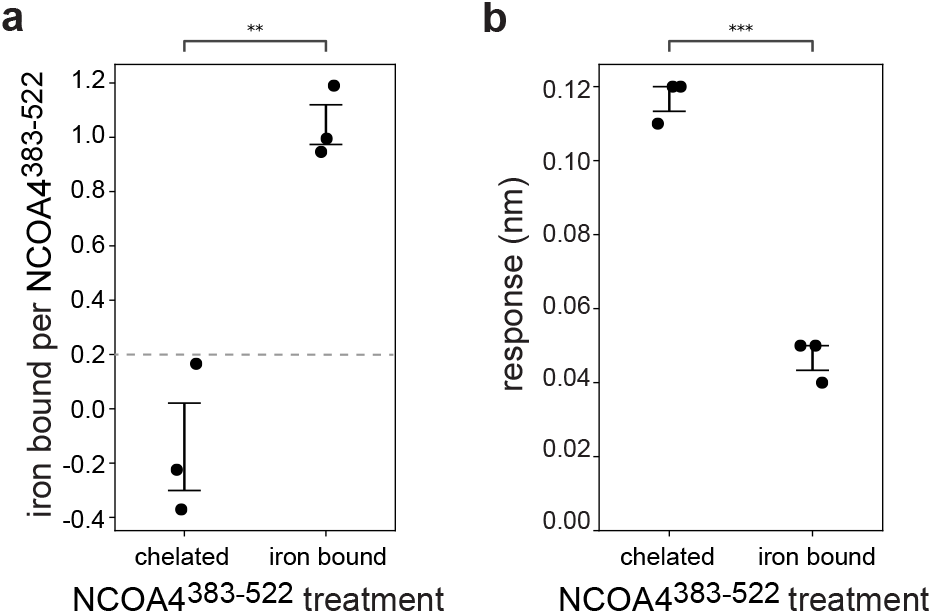
NCOA4^383-522^•(GST-GATE16)_2_ binding is iron dependent. **(a)** Quantification of iron bound per monomer of NCOA4^383-522^ for chelated and reconstituted (iron bound) NCOA4^383-522^ as measured in a ferene assay (Fish, 1988; Levitz et al., 2022; McCarthy & Booker, 2018). Each sample assayed in triplicate, and lower limit of detection noted with dashed line. Statistical significance calculated via independent t-test (ns:p<1, *:p<0.05, **p<0.01, ***p<0.001, ****p<0.0001). **(b)** Quantification of chelated and iron bound NCOA4^383-522^ binding to 10μM (GST-GATE16)_2_ represented as response change in BLI. Each sample assayed in triplicate. Statistical significance calculated via independent t-test as in (a). **(c,d)** Quantification of triplicate ferritin degradation assays described in (a-b). Bars denote standard error of the mean. Fraction ferritin degraded calculated in FIJI (Schindelin et al., 2012) using condition-dependent band intensities as follows:

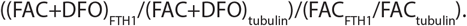 Statistical significance calculated via independent t-test (ns:p<1, *:p<0.05, **p<0.01, ***p<0.001, ****p<0.0001). (**e**)Representative images probing for colocalization of GFP-NCOA4^383-522^ and lysosomes in NCOA4^383-522, 413^F:A, or ^486^F:A cell lines treated with 50μM DFO. Anti-GFP signal shown in green and anti-LAMP1 signal shown in pink. Overlapping areas are white. **(f)** Boxplot of colocalization assay described in (e), with colocalization quantified as a function of area of overlap per nuclei. Black dots represent individual images, with colored dots corresponding to images in (e). Significance calculated via independent t-test (ns:p<1, *:p<0.05, **p<0.01, ***p<0.001, ****p<0.0001). **(g)** Boxplot of colocalization assay described in (e), with colocalization quantified as a function of fraction of lysosomes containing NCOA4^383-522^. Values calculated as Manders’ coefficients in FIJI. Black dots represent individual images, with colored dots corresponding to images in (e). Significance calculated via independent t-test (ns:p<1, *:p<0.05, **p<0.01, ***p<0.001, ****p<0.0001).

## DISCUSSION

Taken together, our biochemical study has revealed that NCOA4^383-522^, the selective autophagy receptor for ferritin, binds directly to GATE16 through two LIR-like motifs:^413^FAECV^417^ and ^485^SFQVI^489^. We found that in isolation each motif binds GATE16 weakly and that NCOA4 relies on avidity between these motifs to achieve tight binding.

Moreover, we observed that to robustly form this complex, GATE16 must be oligomeric, though NCOA4 need not be. Our work further showed that the fragment NCOA4^383-522^ is sufficient for ferritinophagy and that each LIR-like motif is indispensable for this activity *in vivo*. Finally, we found that binding of iron to NCOA4^383-522^ reduces its affinity for GATE16, providing a mechanistic link between cellular iron levels and targeted ferritinophagy.

### Highly avid binding can support LC3/GABARAP-family protein interactions

In cells, LC3/GABARAP-family proteins are relatively abundant (Beck et al., 2011; Huang et al., 2023) and the canonical LIR motif of [W/F/Y]-X-X-[I/L/V] is of low information content and thus common in the human proteome. How then do cells achieve the high binding selectivity one would expect given the degradative capacity of the autophagy-lysosomal pathway? Whereas additional selectivity determinants outside of the core LIR, including a previously described acidic patch N-terminal of the core LIR (Birgisdottir et al., 2013; Johansen & Lamark, 2020), likely play a role, our work additionally suggests that avidly linking multiple LIR-like motifs in a single complex can contribute to specificity and affinity. Under this model selective receptors, or complexes thereof, would need to expose multiple LIRs to support tight binding to LC3s, with such a multivalency requirement adding an additional layer of binding selectivity. Notably, in this model, cells could regulate whether such LIRs were displayed in a condition-specific manner, providing a direct means to modulate selective autophagy.

In support of this avidity model, we have identified two LIR-like motifs in NCOA4^383-522^ that cooperatively facilitate binding to GATE16. In this model, binding of one motif to a single subunit of (GST-GATE16)_2_ increases the local concentration of the dimer-linked GATE16 protomer, which facilitates the second binding event. As we have shown that each motif is indispensable for ferritinophagy *in vivo*, we hypothesize that LC3/GABARAP-family proteins must also act as oligomers to support NCOA4 binding in cells. The fact that the LC3/GABARAPs are conjugated to the autophagosomal membrane, which can proximally situate them at high concentration on a 2D surface, offers a direct mechanism by which they could mimic an oligomeric state. Moreover, whereas our work has focused on GATE16 binding specifically, NCOA4 is thought to interact with multiple members of the LC3/GABARAP family (Mancias et al., 2014), which would allow for avid interactions across the LC3/GABARAP-family members. Notably, our proposed model would also provide a mechanism by which selective autophagy receptors could differentiate between the active, lipid-conjugated forms of LC3/GABARAP-family proteins and their inactive monomeric forms that are found diffusely in the cytosol.

It has become increasingly clear that avidity plays a large role in the progression of autophagy (Zaffagnini & Martens, 2016). In addition to the NCOA4•GATE16 model proposed here, a similar mechanism has been suggested in yeast where avidity is used to support the selective receptor Atg19 binding to Atg8, the yeast homolog of the LC3/GABARAPs (Sawa-Makarska et al., 2014), and in mammalian cells where the selective autophagy receptors p62 and OPTN were each shown to self-oligomerize to facilitate the degradation of their respective cargo (Wurzer et al., 2015; Ying et al., 2010). Moreover, at the most extreme end of low affinity, high avidity interactions, phase separation has been proposed to have a role in autophagic initiation and maturation (Fujioka et al., 2020; Zhang et al., 2018). Our work highlighting the role of avidity in facilitating NCOA4 binding to lipid-conjugated LC3/GABARAPs complements these prior observations.

### An expanded LIR motif

Work over the prior 15 years has resulted in an ever-expanding collection of canonical and non-canonical peptide sequences that support binding to LC3/GABARAP-family proteins (Chatzichristofi et al., 2023; Johansen & Lamark, 2020). One of the peptides we identified (^485^SFQVI^489^) follows the canonical LIR pattern (*i*.*e*., [W/F/Y]-x-x-[I/L/V]), whereas the other (^413^FAECV^417^) is, to our knowledge, the first instance of a cysteine acting as a core hydrophobic residue in a mammalian LIR. By some measures reduced cysteine is strongly hydrophobic and often found in the hydrophobic core of proteins (Nagano et al., 1999), making it a reasonable candidate to occupy one of the hydrophobic pockets on GATE16. Combining this and other observed non-canonical residues at this position (Farnung et al., 2023) with structures demonstrating flexibility in the number of residues separating the two core LIR residues (Keown et al., 2018; Knaevelsrud et al., 2013), leads to a conclusion that the canonical LIR motif is overly stringent, and that the proteome displays an even larger array of potential LC3/GABARAP-family interacting peptides than currently appreciated. Such an expanded understanding of this interface further highlights the challenges in achieving specificity, emphasizing the importance of our described avidity model.

### NCOA4 and ferritin binding

Biochemical (Mancias et al., 2015) and structural (Hoelzgen et al., 2024) studies have pointed to a key role for NCOA4 residues W497 and I489 in ferritin binding. Interestingly, I489 coincides with the final residue of the SFQVI motif that we have shown binds GATE16, which would imply competition between ferritin and GATE16 for NCOA4 binding. Here, we hypothesize that the high-level oligomerization afforded by ferritin’s 432-point symmetry could readily display multiple NCOA4 equivalents per ferritin, and thus enable avid binding through the multiple exposed copies of the FAECV motif. Moreover, the degree of oligomerization could be further expanded in the cellular context, as full-length NCOA4 bears an N-terminal dimerization domain, and others have reported dimerization of the NCOA4^383-522^ fragment (Gryzik et al., 2017). Indeed, Ohshima et al. have observed phase separated condensates composed of NCOA4 oligomers and ferritin that were degraded through autophagy (Ohshima et al, 2022), consistent with the formation of such high-order oligomers. Were such oligomers formed, our model suggests that one equivalent of the ^485^SFQVI^489^ motif could be used to bind GATE16, and the second could associate with ferritin.

### Iron regulation of ferritinophagy

As the process of ferritinophagy releases iron predominantly in response to iron depletion, there are several proposed mechanisms linking ferritinophagy to levels of labile iron. In one model, in iron replete conditions, the E3 ubiquitin ligase HERC2 binds to NCOA4 and promotes its degradation via the proteasome (Mancias et al., 2015; Zhao et al., 2024), thereby negatively regulating ferritinophagy. Additionally, Zhao et al. recently showed that NCOA4 binds an iron-sulfur cluster, which is coordinated using cysteine residues 404, 410, 416 and 422, and that ablation of the iron-sulfur cluster increases NCOA4’s affinity for ferritin (Zhao et al., 2024). Interestingly, cysteine 416 is contained in our described GATE16-binding ^413^FAE<u>C</u>V^417^ motif, raising the possibility of iron-sulfur dependent regulation of this binding activity. Here we showed that in an iron-bound state, NCOA4^383-522^’s affinity for GATE16 is greatly reduced. Hence the presence of iron dually inhibits GATE16 and ferritin binding, and thus ferritinophagy. We postulate that when levels of labile iron fall, this iron-sulfur cluster could be removed from NCOA4, liberating the ^413^FAECV^417^ motif to bind GATE16 and facilitate ferritinophagy, effectively adding an additional layer of iron regulation to the ferritinophagy pathway.

## MATERIALS AND METHODS

### Protein expression and purification

GST-tagged GATE16 was expressed from plasmid pGex-4T-2_GATE-16 (Addgene 73518; pl_JD338) in *E. coli* BL21 Tuner DE3 (st_JD494). His_6_-tagged GATE16 was generated via Q5 site-directed mutagenesis PCR (New England Biolabs), replacing the GST tag with the sequence MHHHHHHGS, and this construct (pl_JD339) was expressed in *E. coli* strain st_JD494. A g-block gene fragment (IDT) containing the sequence for NCOA4^383-522^ was cloned into plasmid pDW363 (Addgene 8842) via HiFi assembly (New England Biolabs) such that it had an N-terminal Avitag followed by a His_6_ tag and a C-terminal MBP (pl_JD337). This vector co-expresses the biotin ligase BirA, enabling NCOA4^383-522^ biotinylation of its Avitag during expression. Mutants of NCOA4^383-522^ (^413^F:A, ^486^F:A, ^413^F:A/^486^F:A, ^413^FAEC:AAAA/^486^F:A, ^413^FAEC:AAAA; corresponding to plasmids pl_JD344 – pl_JD348, respectively) were generated using Q5 site-directed mutagenesis PCR. All NCOA4^383-522^ constructs were expressed in *E. coli* strain st_JD494.

For each protein purified, expression cultures (2L) were grown in 2xYT media at 37°C with aeration, and induced with 1mM isopropyl β-d-1-thiogalactopyranoside at an optical density of ∼0.6. Media for expression of NCOA4^383-522^ constructs was supplemented with 0.05mM biotin. Expression proceeded for three hours at 37°C for GATE16 constructs, or at 30°C for NCOA4^383-522^ constructs before cells were harvested via centrifugation.

Cell pellets bearing GST-GATE16 were resuspended in 50mL buffer PBS (140mM NaCl, 2.7mM KCl, 10mM Na_2_HPO_4_, 1.8mM KH_2_PO_4_, pH 7.3), dounced until homogenous and sonicated using a Qsonica Sonicator (5sec on, 10sec off for 5min, amplitude 35%). The lysate was centrifuged at 251,000g for one hour in a Ti60 rotor, after which the soluble fraction was loaded on a 20mL glutathione-sepharose affinity column (GSTPrep FF 16/10, Cytivia), washed with three column volumes of resuspension buffer, and eluted with 5 column volumes of buffer EB (50mM Tris-HCl, 150mM NaCl, 10mM reduced glutathione, pH 8.0). To generate monomeric GATE16 (mGATE16) for use in fluorescence anisotropy assays, the GST tag was then removed by addition of 20μL thrombin (Millipore 69671-3), and the sample was cleaved at 4°C overnight. The eluate or cleaved fractions for (GST-GATE16)_2_ or mGATE16 respectively, were pooled, concentrated to 2mL and purified over a S75 16/600 size exclusion column in buffer SB (20mM Tris, 150mM NaCl, pH 7.5). Fractions bearing (GST-GATE16)_2_ (∼0.4CVs) or mGATE16 (∼0.7CVs) were identified by SDS-PAGE, pooled, and concentrated via centrifugation in a 30kDa (GST-GATE16) or 3kDa (mGATE16) concentrator (UFC903024 and UFC900324, Millipore Sigma) to a stock concentration of ∼500μM. GST was expressed and purified as described for (GST-GATE16)_2_, using expression plasmid pl_JD340.

Cell pellets bearing His_6_-GATE16 were resuspended in 50mL of buffer NLB (10mM K_2_HPO_4_, 300mM NaCl, 20mM KCl, 10mM imidazole, 5mM 2-mercaptoethanol, pH 8.0), lysed and clarified as above and then loaded onto a 5mL Ni-NTA column (Bio-Rad), washed with two column volumes of buffer NLB, and eluted over 20 column volumes in a linear gradient of buffer NLB with imidazole increasing from 10mM to 1M. Pooled fractions totaling 18mL were then concentrated to 2mL using 3kDa centricon (Millipore Sigma UFC900324) and purified over a S75 16/600 size exclusion column in buffer SB. Fractions bearing His_6_-GATE16 (∼0.7CVs) were identified and concentrated as above.

Cell pellets bearing NCOA4^383-522^ were resuspended in 50mL of buffer NLB, lysed, clarified, and purified on a Ni-NTA column as above. Eluate (∼20mL) was then diluted 5-fold in buffer SB, loaded onto a 5mL MBPTrap HP column (Cytvia), washed with 5 column volumes of buffer SB, and eluted with 8 column volumes of buffer SB supplemented with 10mM maltose. Eluate was pooled, concentrated to 2mL and then purified over a S75 16/600 size exclusion column in buffer SB. Fractions (∼0.4CVs) were identified and concentrated with a 30kDa centricon (Millipore Sigma UFC903024) to ∼50μM.

### Far-Western blots

(GST-GATE16)_2_ at the noted concentrations was spotted (2μL) on a nitrocellulose membrane (Amersham Protran) and allowed to dry at room temperature for 12 minutes. The membrane was then blocked in buffer TBST (20mM Tris, 150mM NaCl, 0.1% Tween-20, pH 7.5) supplemented with 3% bovine serum albumin for 15 minutes at room temperature before being incubated with a 0.1μM solution of the appropriate biotinylated NCOA4^383-522^ protein construct. After washing three times for three minutes with TBST, the membrane was incubated with HRP conjugated streptavidin (Thermo 21130) diluted 1:100,000 and subsequently washed three times in TSBT as above, before applying high sensitivity enhanced chemiluminescent substrate (Thermo 34094). All membranes being compared were imaged concurrently over a two-minute exposure on an Azure Biosystems imager.

### Biolayer interferometry (BLI)

All BLI experiments were conducted on an Octet Red96 instrument (Forte Bio) using streptavidin biosensors (Sartorius 18-5019). Streptavidin biosensors were preincubated with BLI buffer (20mM Tris, 150mM NaCl, 0.05% Tween-20, 1% BSA, 1mM DTT, pH 7.5) for ten minutes. Biotinylated NCOA4^383-522^ constructs were diluted to 50nM in BLI buffer and loaded on the streptavidin biosensors to a response level of 0.6nm. The loaded biosensors were immersed in serial dilutions of the appropriate GATE16 construct at the noted concentrations at an orbital shake speed of 1,000rpm at 30°C for a 250 second association step, with a dissociation step in BLI buffer for 500 seconds between each concentration. For each experiment, background binding of GATE16 to the biosensor was measured using a biosensor lacking NCOA4, and the same process repeated for each concentration. This background binding was subtracted from the resulting assay curves. To generate height response curves as a function of concentration, the response values of the last 50 seconds of each previous dissociation step were averaged and subtracted from the response average of the last 50 seconds of the relevant association step for each concentration.

BLI for chelated and iron bound NCOA4^383-522^ were initiated using protein prepared in a 4°C anaerobic glove box (MBraun, under 100% nitrogen gas) before immediately transfering to the Octet Red96 for assessment. The biotinylated NCOA4^383-522^ constructs were diluted to 250nM in degassed, anaerobic BLI buffer supplemented with 5mM DTT and loaded to a response level of 0.35nm on the streptavidin biosensors. The loaded biosensors were immersed in 10μM (GST-GATE16)_2_ at an orbital shake speed of 1,000rpm at 10°C for a 100 second association step, followed by a dissociation step in anaerobic BLI buffer for 150 seconds. Each measurement was performed in triplicate.

### Spotted peptide array binding assay

A peptide array spanning the NCOA4^383-522^ construct consisted of 20mers offset by three amino acids that were synthesized on a cellulose membrane via SPOT synthesis by the MIT Biopolymers Laboratory. Peptides containing known GATE16 binding peptides (Olsvik et al., 2015; Pankiv et al., 2007; Skytte Rasmussen et al., 2017) derived from p62/SQSTM1 (^332^SGGDDDWTHLSS^343^), ATG4B (^384^EDEDFEILSL^393^) and FYCO1 (^1276^DDAVFDIITDEELCQIQE^1293^) were included as positive controls and a poly His peptide (HHHHHHGSSHHHHHHGSSHH) was added as a negative control. The membrane was briefly soaked in methanol and then washed with TBST before blocking in buffer TBSTB (TBST with 3% BSA) for 30 minutes at room temperature. The membrane was then incubated with 4μg/mL (GST-GATE16)_2_ or GST as a negative control in TBSTB for 30 minutes and subjected to three washes in TBST, each lasting three minutes. Next, bound protein was transferred to a new nitrocellulose membrane at 30V for one hour via wet transfer, and this membrane was blocked in TBSTB for 30 minutes, incubated with HRP conjugated anti-GST antibody (Cytiva RPN1236) at a 1:5,000 dilution, washed with TBST three times for three minutes, exposed to high sensitivity enhanced chemiluminescent substrate (Thermo 34094) and imaged via chemiluminescence (Azure Biosystems Imager).

### Fluorescence anisotropy

Fluorescently labeled (AF Dye-488-TFP ester) and unlabeled peptides were synthesized and HPLC purified by the MIT Biopolymers Laboratory. The peptide sequences used were as follows:

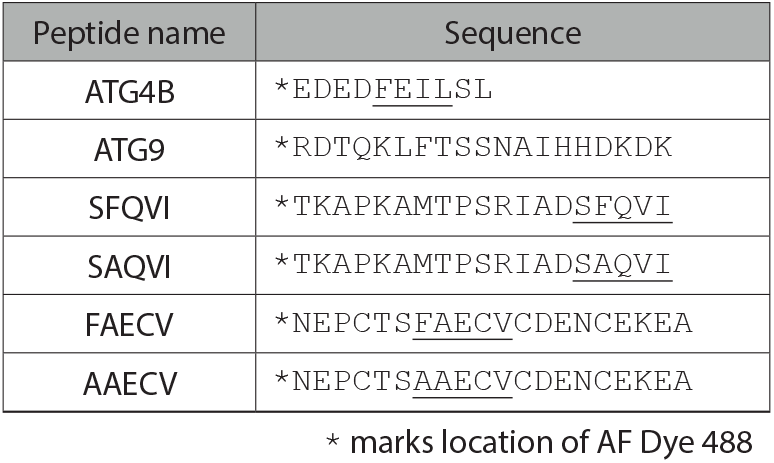

For competitive fluorescence anisotropy assays, 1.5μM mGATE16 was mixed with 10nm labeled ATG4B peptide in a buffer FA (20mM Tris, 150mM NaCl, pH 7.5) at room temperature. Increasing concentrations of the relevant unlabeled peptide were added to individual aliquots of mGATE16 complexed to the labeled ATG4B peptide and incubated for 20 minutes at room temperature. The emission for each 120μL sample at 520nm (excitation at 485nm) for both horizontally (I_VH_) and vertically (I_VV_) polarized signal, along with a G_factor_, was measured on a Photon Technology International (PTI) fluorimeter in a quartz cuvette. From these measurements, an anisotropy value r was calculated by the PTI software as (I_VV_*G_factor_-I_VH_)/(I_VV_+2*G_factor_*I_VH_). ATG9 peptide was used as a negative control as this should not bind to the LDS of GATE16 (Nishimura & Tooze, 2020).

For fluorescence anisotropy assays to measure affinity, 10nM labeled ATG4B peptide was added to individual aliquots of increasing concentrations of His_6_-GATE16 in buffer FA. After the peptide was incubated with His_6_-GATE16 for 20 minutes at room temperature, the horizontally and vertically polarized fluorescence emission at 520nm, along with a G_factor_, was measured as stated above. These same measurements were also taken for all His_6_-GATE16 concentrations as background without labeled peptide. Parallel and perpendicular emission signal intensities for His_6_-GATE16 alone were subtracted from that of His_6_-GATE16 with labeled peptide for each concentration before calculating an anisotropy value r. A curve of anisotropy as a function of His_6_-GATE16 concentration was fit in Graphpad Prism to a binding isotherm to obtain a K_D_.

### Protein thermal stability shift assay

NCOA4^383-522^ constructs were diluted to 10μM in buffer FA and mixed 1:1000 (v/v) SYPRO Orange (Sigma-Aldrich S5692). Each construct was assayed in triplicate in a 384-well plate using an Applied Biosystems QuantStudio 5 Real-Time PCR System. A negative control consisting of SYPRO Orange in buffer was included in all assays. Melt curves for each sample were generated by tracking the fluorescence emission signal at 570nm (excitation at 470nm) as the samples were heated from 25°C to 95°C in 1°C increments. Melting temperatures were calculated as the temperature corresponding to the maximum of the first derivative of the fluorescence signal.

### Size exclusion chromatography coupled to multi-angle light scattering (SEC-MALS)

All SEC-MALS experiments were conducted using a Dawn 8 MALS with an in-line Optilab differential refractive index detector (Wyatt). For both NCOA4^383-522^ and (GST-GATE16)_2_, 100μL of 1mg/mL protein was injected onto an equilibrated WTC-030 HPLC SEC column in buffer FA at a flow rate of 0.5mL/min. The instrument was calibrated with a 2mg/mL BSA standard before each run. Data was analyzed with Astra software (Wyatt) to report apparent molecular weight.

### Western blot assays of ferritinophagy *in vivo*

Stably integrated NCOA4^383-522^ cell lines were generated via retroviral transfection of the relevant NCOA4^383-522^ construct in a CMV-enhancer bearing pMRX plasmid (Addgene 84573) into NCOA4Δ HeLa cells (a gift from Prof. M. Poli, University of Brescia, Italy) (Gryzik et al., 2021). In this pMRX plasmid, EGFP was fused to the N-terminus of NCOA4^383-522^ and its mutants with a 13 residue GS linker. HeLa cells were transfected and selected as described previously (Sena-Esteves & Gao, 2018).

WT HeLa (cl_JD066), NCOA4Δ (cl_JD065), NCOA4^383-522^ (cl_ JD074), ^413^F:A (cl_JD075), and ^486^F:A (cl_JD076) cell lines were seeded in a 12-well cell culture plate in DMEM (Genesee Scientific) and incubated at 37°C, 5% CO_2_. After 24 hours all media was aspirated and replaced with either DMEM or DMEM supplemented with 40μM ferric ammonium citrate (FAC). After an additional 24 hours all media was aspirated and replaced with either DMEM or DMEM supplemented with 50μM deferoxamine (DFO). Following an additional 24-hour incubation, cells were trypsinized (200μL, 5min, 37°C), pelleted via centrifugation at 100g for one minute, resuspended in Dulbecco’s PBS (Sigma Aldrich D8537), pelleted as above before undergoing flash freezing in liquid nitrogen and storage at -80°C.

Frozen cell pellets were resuspended in 100μL 4X Laemmli buffer with Halt protease and phosphatase inhibitor (Thermo Scientific 78440) and lysed with a 29-gauge syringe. Samples were mixed with 285mM dithiothreitol, boiled, and run on an SDS-PAGE gel before being transferred to a PVDF membrane (Invitrogen) using an iBlot2 transfer device (Invitrogen; 7min; 20V). Membranes were blocked overnight in TBSTB, incubated with rabbit anti-FTH1 antibody diluted 1:1,000 (Cell Signaling 3998S) or HRP conjugated anti-tubulin antibody diluted 1:20,000 (GeneTex GTX628802-01) as above. Anti-rabbit HRP conjugated secondary antibody (ABclonal AS014) was applied at a 1:1,000 dilution and washed, exposed to enhanced chemiluminescent substate, and imaged as above. Ferritin levels were then measured by densitometry in FIJI (Schindelin et al, 2012) and normalized to tubulin intensity measured as above.

### Fluorescence co-localization assays

NCOA4^383-522^ cell lines were seeded on glass coverslips in a 12-well cell culture plate in DMEM as above. After 24 hours, media was aspirated and replaced with DMEM supplemented with 50μM DFO, and cells were incubated an additional 24 hours. Cells were then fixed with 4% paraformaldehyde in PBS (150mM NaCl, 2.7mM KCl, 10mM Na_2_HPO_4_, 1.8mM KH_2_PO_4_, pH 8.0) for 15 minutes at room temperature, quenched with 100mM glycine in PBS for 10 minutes, and washed three times with PBS. Cells were next permeabilized with 0.5% Triton X-100 in PBS for 10 minutes, washed once with PBS and blocked for 30 minutes in 2% BSA in PBS. Slides were next incubated in anti-GFP mouse antibody (Santa Cruz sc-9996) diluted 1:200, anti-LAMP1 rabbit antibody (Cell Signaling 9091T) diluted 1:200, and Hoechst stain (Invitrogen H3570) diluted 1:1,000, in 2% BSA in PBS for one hour. Following three 5-minute soaks in PBS, slides were incubated in Alexa Fluor 488 anti-mouse antibody (Thermo Fisher A-11001) diluted 1:200, and Alexa Fluor 568 anti-rabbit antibody (Thermo Fisher A-11036) diluted 1:200, for one hour. Slides were then soaked three times for 5 minutes in PBS, dried and mounted on slides with ProLong Gold mounting medium, set overnight, and sealed. Slides were imaged on a Dragonfly 505 spinning-disk confocal microscope with an iXon Ultra 888 EMCCD camera with the following settings: 60x objective pixel size (206µm*206µm), 40µm pin hole size, 0.5µm interval z stack slices, 405nm laser with 445/46 bandpass emission filter, 488nm laser with 521/38 bandpass filter and 561nm laser with 594/43 bandpass filter. A minimum of 100 cells were imaged per condition.

All image analysis was conducted in FIJI (Schindelin et al., 2012). To measure the area of overlap per cell, the best focused image from the z-stack was first manually chosen based on the nuclei in the 405nm channel. The in-focus image was then split into its respective green (NCOA4^383-522^) and pink (lysosome) channels before merging to a single RGB image. Images were viewed at fixed brightness and saturation when selecting regions of overlap (white). The overlapping regions were then measured and the area per nuclei recorded. The JACoP plugin for FIJI (Bolte & Cordelières, 2006) was used to estimate the fraction of lysosomes that colocalized with NCOA4^383-522^. In brief, each z stack was split into the respective channels, background subtracted with a rolling ball radius of 50 pixels and set to standard brightness values. For all images, threshold values were manually chosen for the pink and green channels, and the Manders’ value, which calculates the fraction of pink signal that overlaps green signal was recorded.

### NCOA4^383-522^ iron chelation and reconstitution

To obtain reconstituted NCOA4^383-522^ sample, purified NCOA4^383-522^ was diluted two-fold in buffer IC (20mM Tris, 150mM NaCl, 5mM DTT, pH 7.5, degassed and stored anaerobically) incubated with a 10-fold molar excess of ammonium iron (II) sulfate hexahydrate and sodium sulfide nonahydrate overnight at 20°C in an anaerobic glove box. Excess iron and sulfur were pelleted via centrifugation at 17,200g for 2mins and filtered using a centrifuge tube 0.22μm filter (Corning 8160) before loading onto an S200 10/300 increase size exclusion column run in buffer IC under anaerobic conditions. Fractions eluting at ∼0.57 column volumes were collected and concentrated with a 30kDa centricon for use in BLRI.

To obtain chelated sample, purified NCOA4^383-522^ was incubated with EDTA and potassium ferricyanide in molar ratios of 1:50 and 1:20 respectively for 30min at room temperature. This material was run over an S200 10/300 increase in buffer IC. Fractions eluting at ∼0.57 column volumes were collected and concentrated for subsequent use in BLI.

Chelated and reconstituted NCOA4^383-522^ were both tested in a ferene assay (Fish, 1988; Levitz et al., 2022; McCarthy & Booker, 2018) to measure iron concentration. In brief, chelated and reconstituted NCOA4^383-522^ samples were diluted to 2.5μM in buffer IC and iron (III) nitrate nonahydrate (Thermo 047282.AP) standards spanning 0 to 100μM were made in volumes of 100μL. Samples and standards were mixed with 100μL of Reagent A (156mM SDS, 113mM saturated sodium acetate) and 100μL Reagent B (274mM ascorbic acid, 8mM sodium meta-bisulfite, 536mM saturated sodium acetate) then incubated at 30°C for 15 minutes. 5μL of Reagent C (36mM ferene) was added before centrifuging samples at 21,130g for 5min. The absorbance at 592nm for 250μL of supernatant from each condition was then measured in a 96-well clear plate using a Molecular Devices plate reader. Three replicates of each protein sample were compared to a standard curve calculated from the standards to obtain an iron concentration.

## ACKNOWLEDGEMENTS

We thank Jennifer Kosmatka, Dante Avalos, Cathy Drennan, Michael Das, and Allen Sanderlin for helpful discussion, and the MIT Biophysical Instrumentation Facility and MIT Biopolymer Laboratory for supporting access to key instrumentation. This work was supported by NIH grants 5T32-GM007287, and NSF-CAREER grant 2046778, and the Whitehead Family.

## AUTHOR CONTRIBUTIONS

AL and JHD conceived the work, designed and interpreted the experiments, and wrote the manuscript. AL performed all experiments. The authors declare that they have no conflicts of interest.

## SUPPLEMENTARY INFORMATION

**Supplementary Figure 1.**
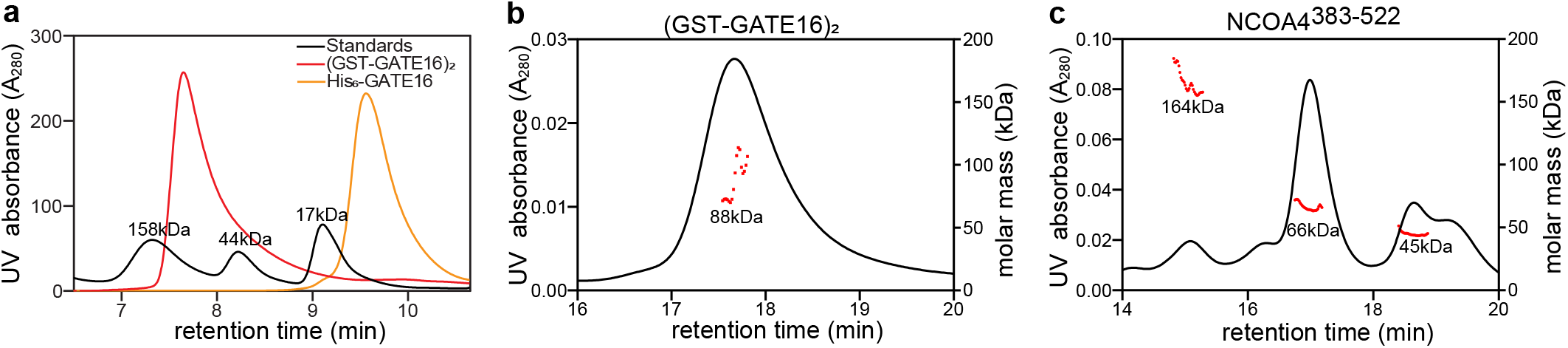
Oligomeric states of GATE16 and NCOA4^383-522^ in solution. Related to Figure 1. **(a)** HPLC trace of (GST-GATE16)_2_ and His_6_-GATE16. Gel filtration standards annotated with molecular weight. Note a monomer of His_6_-GATE16 is 15kDa in mass. **(b)** SEC-MALS analysis of (GST-GATE16)_2_. Black curve traces UV absorbance at 280nm. Red dots represent determined molar mass across the given peak. Measured mass of peak is 88±14kDa. Note a dimer of GST-GATE16 is 80kDa in mass. **(c)** SEC-MALS analysis of NCOA4^383-522^. Black curve and red dots as in (b). Measured masses from peaks left to right are 164±7.3kDa, 66.2±2.0kDa, 44.8±2.0kDa. Note a monomer of the biotinylated AviTag-His_6_-NCOA4^383-522^-MBP construct used in all *in vitro* assays is 61kDa and isolated MBP is 40kDa in mass.

**Supplementary Figure 2.**
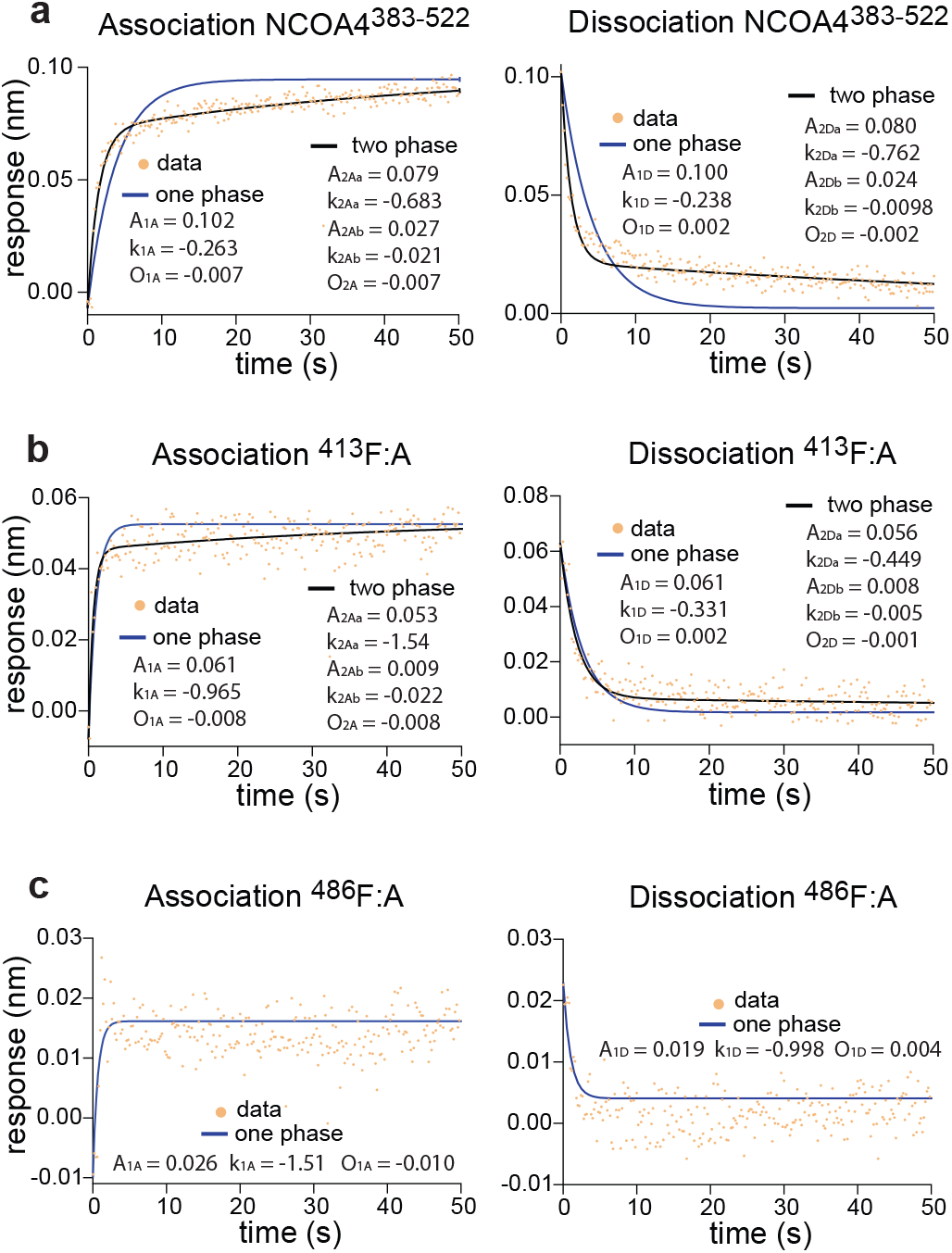
NCOA4^383-522^ and (GST-GATE16)_2_ binding kinetics measured by BLI. Related to Figure 1. **(a)**BLI association and dissociation curves of NCOA4^383-522^ binding to 5μM (GST-GATE16)_2_. BLI response data fit in Prism to one- and two-phase models as follows:

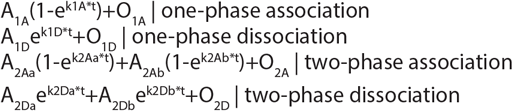 **(b)**BLI reponse curves of ^486^F:A binding fit as in (a). **(c)**BLI response curves of ^413^F:A binding fit as in (a).

**Supplementary Figure 3.**
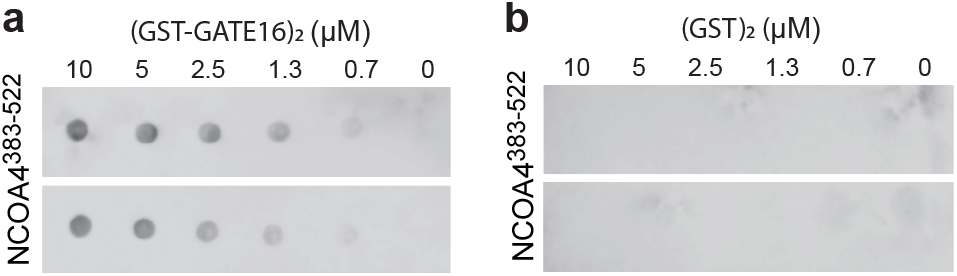
Assessment of nonspecific binding of NCOA4^383-522^ to GST in far-westerns. Related to Figure 1. **(a)**Far-western of NCOA4^383-522^ binding to (GST-GATE16)_2_, which was spotted on membrane at indicated concentration. NCOA4^383-522^ incubated on membrane at 0.1μM. Signal detected by HRP conjugated streptavidin. Two replicates shown. **(b)**Far-western as in (a), with GST replacing (GST-GATE16)_2_.

**Supplementary Figure 4.**
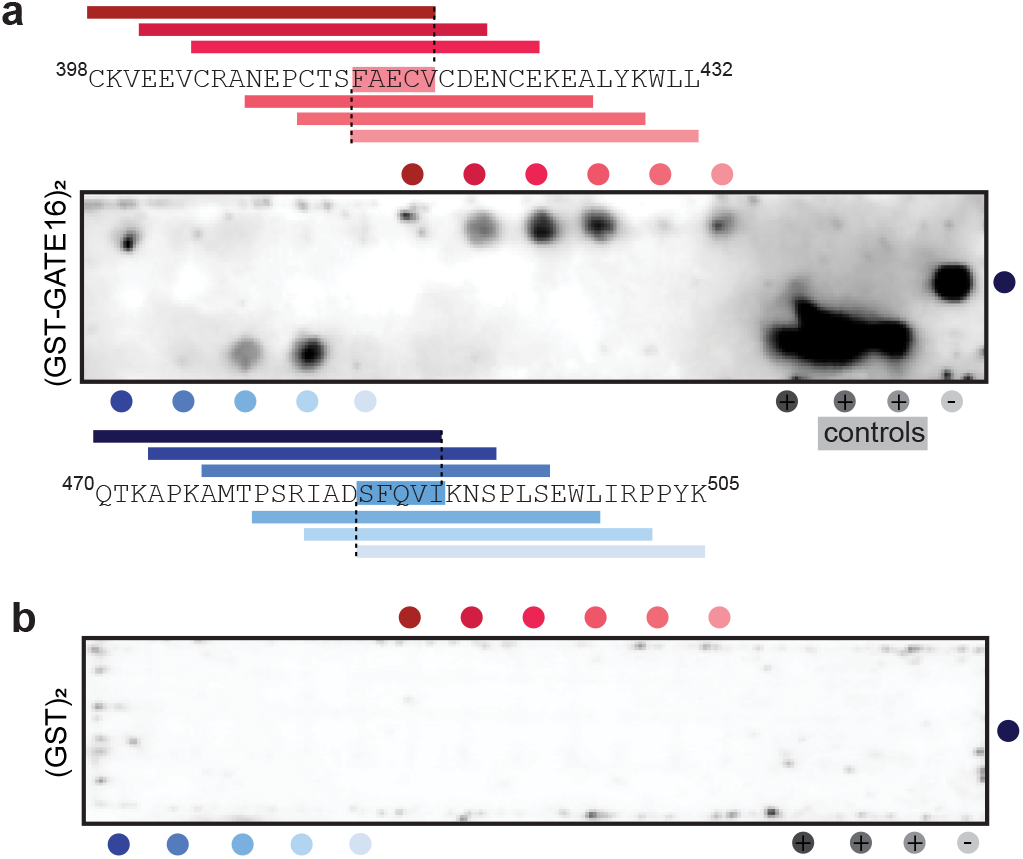
NCOA4^383-522^ binding to (GST-GATE16)_2_ via peptide array. Related to Figure 2. **(a)**Binding of (GST-GATE16)_2_ to a peptide spot array composed of 20mers tiled across the NCOA4^383-522^ sequence. (GST-GATE16)_2_ binding detected using an HRP-conjugated anti-GST antibody. Sites of (GST-GATE16)_2_ binding are indicated by colored dots adjacent to peptide array, with corresponding peptide sequence spans depicted. Positive control peptides arrayed left-to-right are noted with grey circles, and were derived from p62/SQSTM (SGGDDDWTHLSS), ATG4B (EDED<u>FEIL</u>SL), and FYCO1 (DDAVFDIITDEELCQIQE), with known LIR motif underlined. Negative control peptide noted by light grey circle corresponded to a poly-His peptide (HHHHHHGSSHHHHHHGSSHH). **(b)**Far-western as in (a), with GST replacing (GST-GATE16)_2_.

**Supplementary Figure 5.**
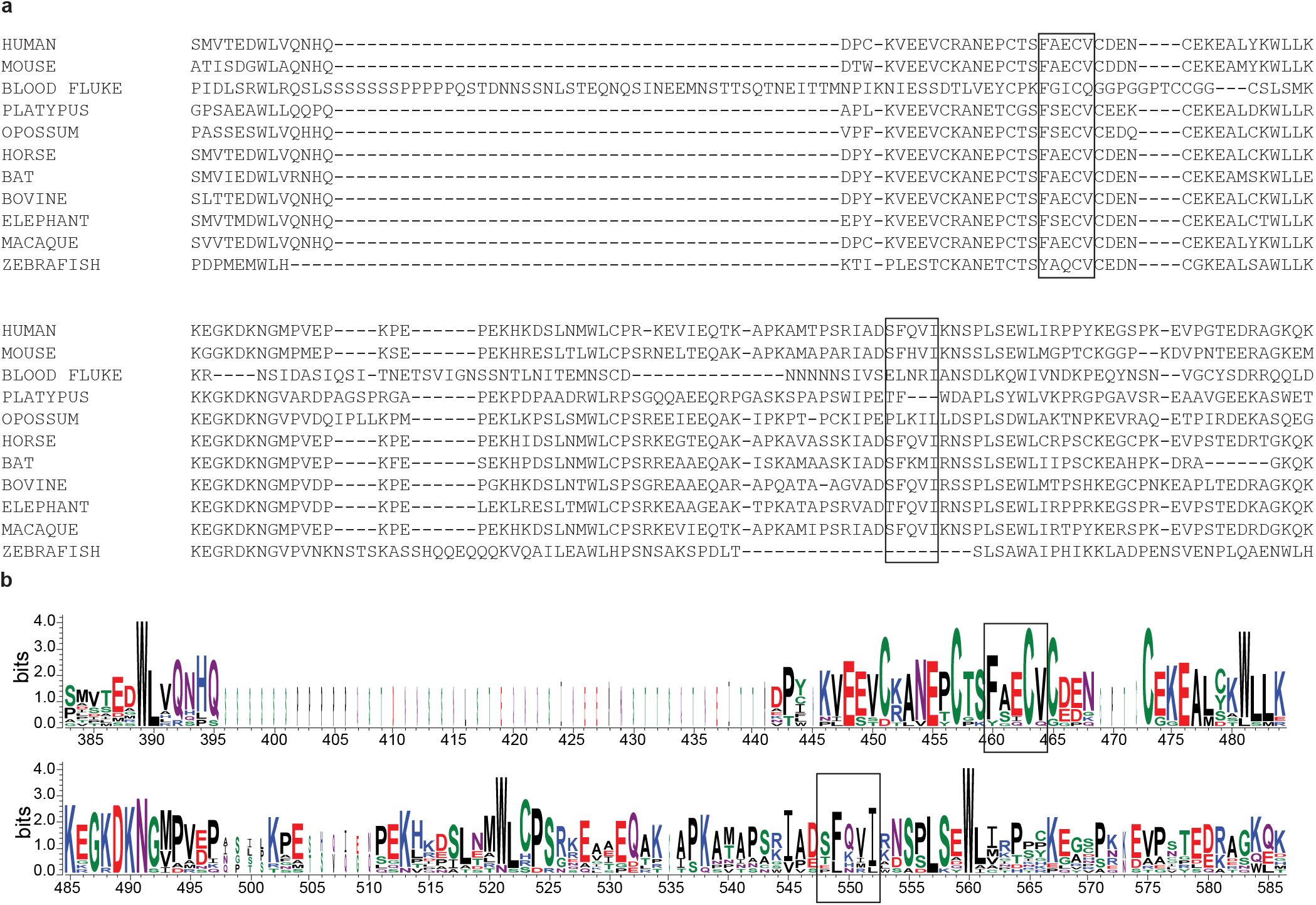
Sequence conservation of the two LIR-like motifs of NCOA4^383-522^. Related to Figure 2. **(a)**Sequence alignment of human NCOA4^383-522^ and orthologs. Identified LIR-like motifs are indicated with boxes. **(b)**Sequence logo of the alignment in (a) generated by WebLogo 3.7.4 (Crooks et al., 2004). Motifs are indicated with boxes. Height at each position indicates overall sequence conservation at that position and height of individual letters at each position indicates relative frequency of the residue for that position. The width of each position is proportional to the number of aligned sequences at that position.

**Supplementary Figure 6.**
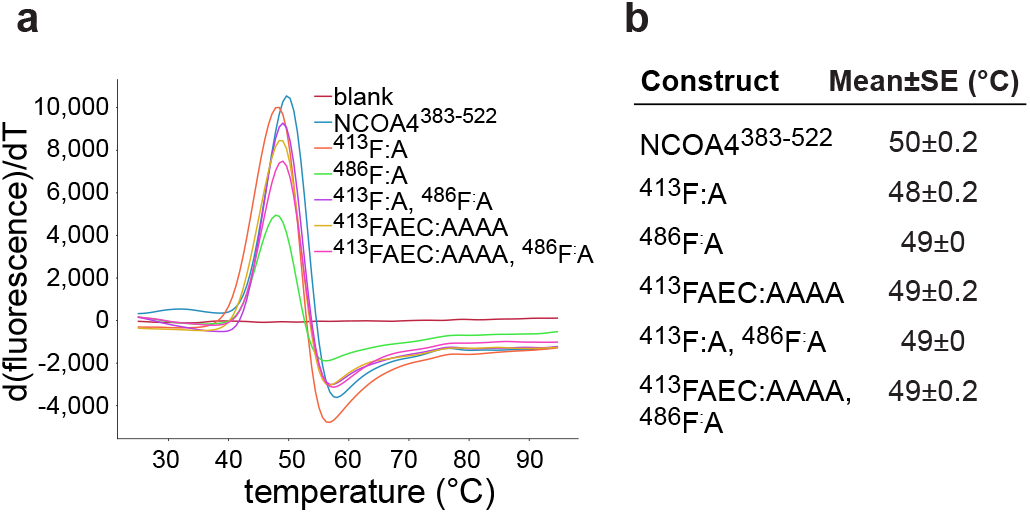
Impact of NCOA4^383-522^ mutants on protein stability. Related to Figure 2. **(a)**Protein thermal shift assay melting curves plotting the first derivative of SYPRO Orange fluorescence emission as a function of temperature for NCOA4^383-522^ and NCOA4^383-522^ mutants (see Methods). **(b)**Table of melting temperature for NCOA4^383-522^ and NCOA4^383-522^ mutants measured in (a). Mean and standard error of three replicates for each construct denoted.

**Supplementary Figure 7.**
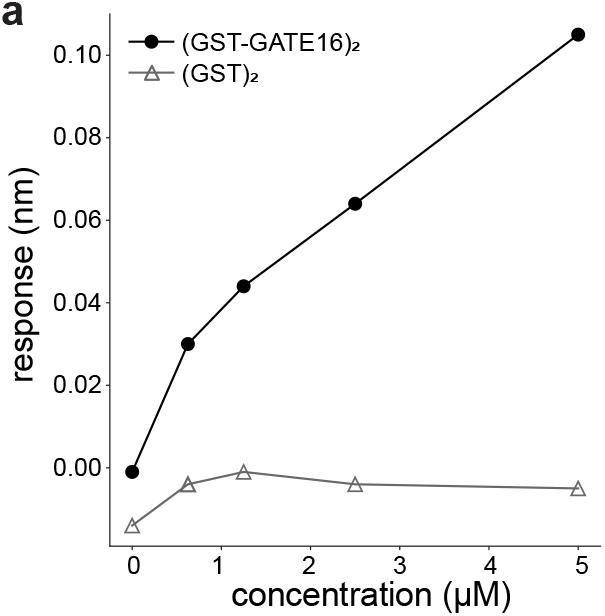
Assessing nonspecific association of NCOA4^383-522^ to (GST)_2_. Related to Figure 3. **(a)**BLI response measurement of NCOA4^383-522^ binding to (GST-GATE16)_2_ and isolated (GST)_2_ alone where NCOA4^383-522^ construct was immobilized to the probe and tested against increasing concentrations (0nM, 625nM, 1.25μM, 2.5μM, 5μM) of either GST construct. Response height measured at each concentration is plotted. time (s)

**Supplementary Figure 8.**
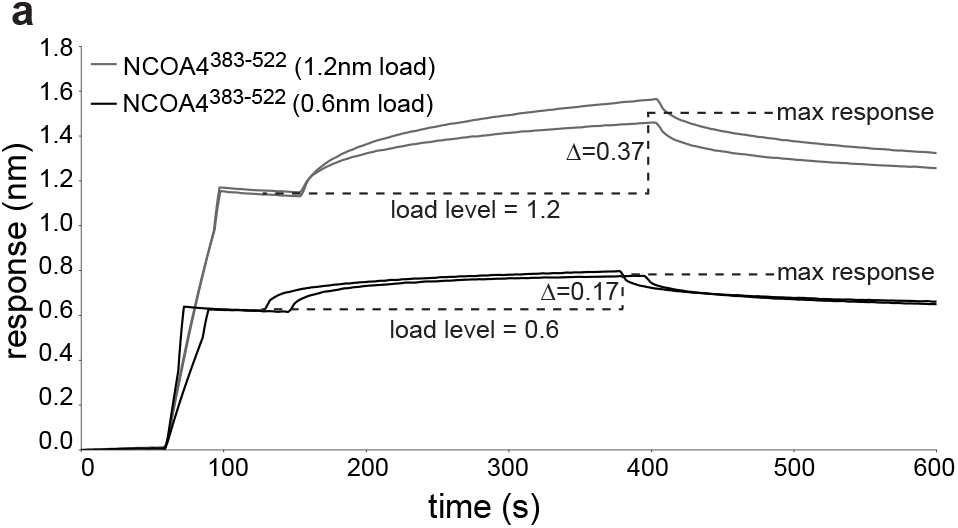
Load-dependent BLI response. Related to Figure 3. **(a)**BLI association and dissociation curves of NCOA4^383-522^ binding to 5μM (GST-GATE16)_2_ where NCOA4^383-522^ construct is immobilized to the probe. Replicates at NCOA4^383-522^ loading levels of 0.6 nm and 1.2 nm shown. Average BLI response across replicates denoted as difference (Δ) of max response and load level.

**Supplementary Figure 9.**
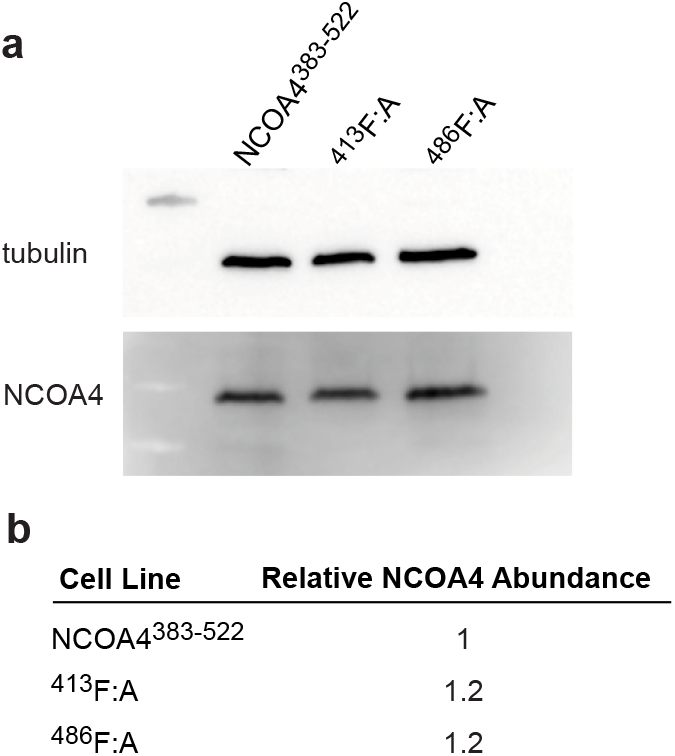
Cellular levels of NCOA4^383-522^ and associated mutants. Related to Figure 4. **(a)**Western blot assessing levels of NCOA4^383-522, 413^F:A, and ^486^F:A expressed under steady-state conditions in HeLa cell lines. NCOA4 levels probed using an anti-GFP antibody against the GFP-fused NCOA4. Tubulin levels also probed by Western to assess relative sample loading. **(b)**Table reporting NCOA4 abundance of each mutant relative to NCOA4^383-522^ as measured in (a). In each instance, levels were normalized using the tubulin loading control.

## Notes

### Competing Interest Statement

The authors have declared no competing interest.

